# TRIC-Based High-Throughput Screening Enables the Discovery of Small Molecule CD28 Binders

**DOI:** 10.1101/2025.05.27.656499

**Authors:** Laura Calvo-Barreiro, Moustafa T. Gabr

## Abstract

CD28 is a pivotal costimulatory receptor involved in T cell activation and immune regulation, positioning it as a key therapeutic target for inflammatory diseases, including inflammatory bowel disease (IBD). Despite its potential, small molecules targeting CD28 are still limited. To fill this gap, we developed a high-throughput screening (HTS) platform based on Temperature-Related Intensity Change (TRIC) technology, enabling rapid, immobilization-free screening of chemical libraries of small molecules. Using the Dianthus instrument, we applied our optimized TRIC assay for CD28 (signal-to-noise ratio of 21.99) to screen two MedChemExpress libraries: Small Molecule Immuno-Oncology Compounds (SMIOC) and Protein-Protein Interaction Inhibitors (PPII), identifying 50 initial hits. Following exclusion of compounds with dye interference or aggregation artifacts, 12 candidates were prioritized for further validation. Microscale thermophoresis (MST) confirmed dose-dependent binding of seven compounds to CD28, with affinities in the micromolar range. Surface plasmon resonance (SPR) further validated two compounds, EABP 02303 and CTEP, as CD28 binders. These results demonstrate that our TRIC-based HTS platform is robust, scalable, and effective for identifying small molecule CD28 binders. The incorporation of orthogonal validation supports the reliability of our findings and highlights the feasibility of small-molecule discovery targeting CD28.

## 1. Introduction

Cluster of Differentiation 28 (CD28) and Inducible T Cell Costimulator (ICOS) are closely related members of the CD28 family of costimulatory receptors, which play pivotal roles in T cell activation, survival, and differentiation. Both are transmembrane glycoproteins expressed on T cells and function by engaging ligands on antigen-presenting cells to modulate immune responses. While CD28 is constitutively expressed and essential for the initiation of T cell activation, ICOS is induced upon activation and contributes to T cell proliferation and cytokine secretion. CD28 binds to CD80 (B7-1) and CD86 (B7-2) on antigen-presenting cells, initiating downstream signaling cascades such as PI3K/Akt and NF-κB that are critical for full T cell activation and survival^1-3^. Owing to the complementary and partially overlapping functions of CD28 and ICOS, these receptors have been implicated in the pathogenesis of several immune-mediated disorders, including inflammatory bowel disease (IBD)^1-3^.

To date, dual inhibition strategies of CD28 and ICOS have primarily relied on Fc fusion protein-based biologics^4-6^. However, such protein therapeutics are often associated with challenges including immunogenicity, anti-drug antibody formation, and increased risk of adverse effects^7^. In contrast, small molecules offer significant advantages, including reduced immunogenic potential and improved pharmacokinetic flexibility. Our previous efforts led to the discovery of small molecule and peptide inhibitors of ICOS, including modulators of the ICOS/ICOS-L protein– protein interaction^8-11^, however, remains a substantial gap in the availability of small molecules capable of targeting CD28.

To address this unmet need and enable high-throughput screening (HTS) of chemical libraries of small molecules for affinity binding to CD28 on a large scale, we developed a primary screening method based on Temperature-Related Intensity Change (TRIC) technology. This method fully exploits the capabilities of the Dianthus instrument (NanoTemper Technologies), chosen for its mass-independent, immobilization-free, and plate-based detection advantages.

TRIC technology is based on the detection of fluorescence intensity changes induced by a controlled temperature increase. When a target protein is labeled with a fluorescent dye, its local microenvironment can influence the brightness of the emitted light—especially when a small molecule ligand binds to the protein, causing structural or conformational changes. These fluorescence variations, often subtle at constant temperature, become more pronounced with a slight temperature increase. TRIC leverages this principle by applying a rapid, localized heating step using an infrared laser to enhance the sensitivity of detection and enabling robust discrimination between bound and unbound states.

This immobilization-free, solution-based detection method offers a valuable alternative to conventional biophysical assays, particularly for challenging targets like immune checkpoint receptors. Our TRIC-based platform addresses a critical gap in the discovery of small molecule immunomodulators of CD28—a key costimulatory receptor with limited chemical tractability to date. By enabling efficient identification of CD28-binding compounds, our approach complements ongoing efforts in ICOS-targeted drug development and supports the advancement of dual-target therapeutic strategies for IBD and related immune disorders.

## 2. Methods

### 2.1. Screened Libraries

Two libraries from MedChemExpress (MCE) were chosen as candidate libraries for the validation of the designed workflow: Small Molecule Immuno-Oncology Compound Library (Cat# HY-L031, MCE, Monmouth Junction, NJ, USA) and Protein-Protein Interaction Inhibitor Library (Cat# HY-L109, MCE). All screened compounds were contained in 384-well plates and dissolved in DMSO at a concentration equal to 10 mM. Libraries were stored at -80°C upon arrival and until use.

After HTS of stock plates, new lyophilized material for every selected small molecule candidate was acquired from MCE, reconstituted in DMSO at a 50 mM stock concentration and stored at -30°C until use.

### 2.2. Dianthus: High-Throughput Affinity Screening Platform

The Monolith His-Tag Labeling Kit RED-tris-NTA 2^nd^ Generation (Cat # MO-L018, NanoTemper Technologies, München, Germany) was used for the labeling of the human CD28 Protein, His Tag (Cat # CD8-H52H3, Acro Biosystems, Newark, DE, USA) following manufacturers’ instructions. After assessing the affinity of the His-labeling dye for the His-tagged human CD28 protein, a solution containing 20 nM of the RED-tris-NTA dye and 40 nM of the human CD28 protein was prepared in assay buffer. The mixture was incubated in the dark at room temperature for 30 minutes prior to each experiment.

The labeled protein was incubated at a 1:1 ratio with the corresponding compound or control for 15 minutes at room temperature in the dark. Small-molecule compounds were screened at a final concentration of 200 μM in assay buffer containing 2% DMSO. Human CD80, Fc Tag (Cat. #B71-H5259, Acro Biosystems), at 2 μM in assay buffer supplemented with 2% DMSO, was used as the positive control, while assay buffer with 2% DMSO alone served as the negative control.

To ensure proper solution homogenization, the mixture was pipetted 15 times. After incubation, the assay plate was centrifuged at 1,000 × g for one minute to eliminate any bubbles in the wells. Following centrifugation, the sample was loaded into the Dianthus NT.23 Pico (NanoTemper Technologies). The assay buffer consisted of 1x PBS with 0.005% Tween-20 (pH=7.4) and the reaction volume was 20 μL for all experiments.

The equipment was set to a temperature of 25 °C. Fluorescence intensity at 670 nm was measured for 1 second without heating, followed by 5 seconds with the infrared laser activated for each sample (TRIC trace). The results are presented as normalized fluorescence (F_norm_, ‰), calculated as the ratio of fluorescence after 5 seconds of infrared laser activation (F_hot_) to the fluorescence before activation (F_cold_). All TRIC traces were normalized to start at 1 (relative fluorescence).

In the HTS experiment, one technical replicate was performed per compound, while 32 technical replicates were conducted for the negative control at regular intervals in the 384-well plate. Negative control wells were strategically positioned to evaluate the laser performance of the Dianthus NT.23 Pico instrument, which utilizes two lasers—one reading the top half and the other screening the bottom half of the assay plate. Additionally, these controls ensured that protein behavior remained consistent throughout the reading run, which lasted approximately 35 minutes.

Once excluding the compounds that interacted with the RED-tris-NTA dye alone, binding affinity experiments were performed using the Monolith NT.115 (Nanotemper Technologies). Data was acquired and analyzed using DI.Control and DI.Screening Analysis software (NanoTemper Technologies), respectively.

After applying the selection criteria (*details in the Results and Discussion section*), candidate small molecules were identified as potential binders to the human CD28 protein. Two additional technical replicates were performed to confirm binding. If binding was confirmed, a control experiment was conducted for each selected small molecule to ensure that none of the compounds interfered with the RED-tris-NTA dye alone—either by quenching its signal, increasing its fluorescence intensity, or altering the TRIC trace. Compounds that exhibited any interaction with the dye alone were excluded from further analysis. Subsequently, binding affinity experiments were performed using the Monolith NT.115 (NanoTemper Technologies). Data acquisition and analysis were carried out using DI.Control and DI.Screening Analysis software (NanoTemper Technologies), respectively.

### 2.3. Monolith: Binding Affinity Screening Platform

As in the HTS using the Dianthus NT.23 Pico (NanoTemper Technologies), the Monolith His-Tag Labeling Kit RED-tris-NTA 2^nd^ Generation (NanoTemper Technologies) was used to label the His-tagged human CD28 protein (Acro Biosystems), following the manufacturer’s instructions. After confirming the affinity of the RED-tris-NTA dye for the His-tagged CD28 protein, a solution containing 50 nM of the dye and 100 nM of the protein was prepared in assay buffer. The mixture was incubated in the dark at room temperature for 30 minutes prior to each experiment.

Sixteen-point serial dilutions were prepared for each candidate small molecule in assay buffer to the corresponding 2x final concentration in the presence of 4% DMSO. The assay buffer consisted of 1x PBS with 0.005% Tween-20 (pH=7.4) and the reaction volume was 15 μL for all experiments. After, the labeled protein was incubated at a 1:1 ratio with the corresponding compound for 15 minutes at room temperature in the dark. To ensure proper solution homogenization, the mixture was pipetted 15 times. After incubation, each sample was dipped into a Monolith Premium Capillary (Cat# MO-K025, NanoTemper Technologies) and loaded into the Monolith NT.115 instrument (NanoTemper Technologies).

The equipment was set to a temperature of 25 °C. Samples were measured for 1 second without heating (before infrared laser activation) and during 20 seconds with the infrared laser turned on. The Expert Mode on the MO.Control Software (NanoTemper Technologies) was used to obtain experimental data using 80% excitation power and MST laser at medium power for every screened compound. Results are displayed as fraction bound and plotted against ligand concentration. The fraction bound is calculated by dividing each ΔF_norm_ value by the curve amplitude, resulting in a normalized value between 0 and 1 for each data point.

Three independent experiments were performed to determine the affinity constants. Data was acquired and analyzed using MO.Control and MO.Affinity Analysis software (NanoTemper Technologies), respectively.

Compounds that demonstrated consistent affinity constants in the binding affinity experiments were further selected for confirmatory testing using orthogonal validation by surface plasmon resonance (SPR).

### 2.4. Surface Plasmon Resonance: Orthogonal Validation Platform

Binding analysis was performed using a Biacore™ 8K instrument (Cytiva, Marlborough, MA, USA) at 25 °C with 1x PBS-P+ buffer (Cat# 28995084, Cytiva) supplemented with 2% DMSO as the running buffer. Biotinylated human CD28 protein (His-tagged, AviTag™; Cat# CD8-H82E5, Acro Biosystems) was captured on a Series S Sensor Chip CAP at a concentration of 50 μg/mL, using a flow rate of 10 μL/min for 300 seconds. This resulted in an immobilization level of approximately 1,750 RU, using the Biotin CAPture Reagent (included in the Biotin CAPture Kit, Series S; Cat# 28920234, Cytiva), which was diluted 1:1 with 1x HBS-EP buffer (Cat# BR100826, Cytiva). Besides the modified streptavidin-based Biotin CAPture Reagent, the Biotin CAPture Kit includes two regeneration solutions. When combined and applied at the end of the experiment, these solutions remove the Biotin CAPture Reagent, the ligand, and any bound analyte from the chip surface, allowing for chip reuse.

For binding analysis, a multi-cycle kinetics and affinity approach was established. Serial dilutions of the analytes (ranging from 500 μM to 3.91 μM), along with a 0 μM reference, were prepared in running buffer and injected over the chip at a flow rate of 30 μL/min for 90 seconds (association phase), followed by a 240-second dissociation phase. An anti-CD28 antibody (Cat# K123A, Promega, Madison, WI, USA) at 2 μg/mL was used as a positive control in each channel to confirm protein activity at the end of the experiment.

Prior to loading plates into the instrument, plates were centrifuged at 1,000 rpm for one minute at room temperature and sealed with the recommended sealing film (Cat# BR100577 and Cat# 28975816, Cytiva). A solvent correction run was also performed before and after the positive control step. Data were acquired using Biacore 8K Control Software (Cytiva) and analyzed by non-linear curve fitting using steady-state affinity analysis in Biacore™ Insight Evaluation Software (Cytiva).

## 3. Results and Discussion

### 3.1. Development of an Affinity-Based Screening Assay and Workflow Using TRIC Technology

The extracellular region of the His-tagged human CD28 protein (Asn19–Pro152, accession #P10747-1) was selected as the target for labeling based on the following criteria: i) our interest lies in small molecules that bind to the extracellular portion of the human protein; ii) it forms an active, glycosylated, disulfide-linked homodimer that mimics the physiologically relevant structure found in humans; and iii) the polyhistidine tag at the C-terminus is unlikely to interfere with the protein’s structure or function due to its position and relatively small size compared to other potential tags. Based on this, we employed a His-Tag labeling strategy for our target protein. After verifying that the protein-dye binding affinity was below 10 nM in all tested assay buffers (*composition detailed below*), we optimized the buffer condition (Table 1) for the subsequent experiments. Human CD80 protein at 2 μM [corresponding to the reported dissociation constant (K_d_) for its interaction with human CD28 as determined by SPR on a Biacore T200 system (K_d_ =2.01 μM; ACROBiosystems, Cat. No. B71-H5259)^12^] served as the positive control to identify the assay buffer that maximized the signal difference between the bound and unbound states. The following assay buffers were evaluated: i) 1x PBS supplemented with 0.005% Tween-20 (pH=7.4); ii) 10mM HEPES, 150 mM NaCl, supplemented with 0.005% Tween-20 (pH=7.4); and iii) 50mM HEPES, 150 mM NaCl, supplemented with 0.005% Tween-20 (pH=7.4). As shown in Table 1, the PBS-based buffer demonstrated superior performance, with a ΔF_norm_ of 22.53 and a signal-to-noise ratio (SNR) of 21.99, compared to the 10 mM HEPES buffer (ΔF_norm_ = 14.25, SNR = 6.97) and the 50 mM HEPES buffer (ΔF_norm_ = 19.51, SNR = 8.37) in detecting binding with the positive control. Consequently, the PBS-based buffer was selected for subsequent screening.

**Table 1.** Buffer Optimization of the CD28 TRIC assay.

| Buffer composition | pH | Signal-to-noise ratio | $\Delta F_{\text{norm}}$ |
| --- | --- | --- | --- |
| 1x PBS, 0.005% Tween-20 | 7.4 | 21.99 | 22.53 |
| 10 mM HEPES, 150 mM NaCl, 0.005% Tween-20 | 7.4 | 6.97 | 14.25 |
| 50 mM HEPES, 150 mM NaCl, 0.005% Tween-20 | 7.4 | 8.37 | 19.51 |

Two libraries from MedChemExpress (MCE) were chosen as suitable candidate libraries for the validation of the designed workflow: Small Molecule Immuno-Oncology Compound (SMIOC) Library and Protein-protein Interaction Inhibitor (PPII) Library (**Figure 1**). The SMIOC Library is a high purity and quality 493-member small molecule library (validated by NMR and LC/MS) that holds bioactive tumor immunology compounds that target some important checkpoints, such as CTLA-4 and PD-1/PD-L1, among others. Thus, since CD28 belongs to a subfamily of costimulatory molecules that are characterized by an extracellular variable immunoglobulin-like domain that also include homologous receptors such as ICOS, CTLA-4, PD-1, among others^13^; we thought that this library might be a good fit for CD28-targeted assay development. Likewise, the PPII Library composed by 569 compounds was chosen based on our goal of targeting PPI, such as CD28/CD80 and CD28/CD86, as a new direction in treating diseases and an essential strategy for the development of new drugs.

**Figure 1.**
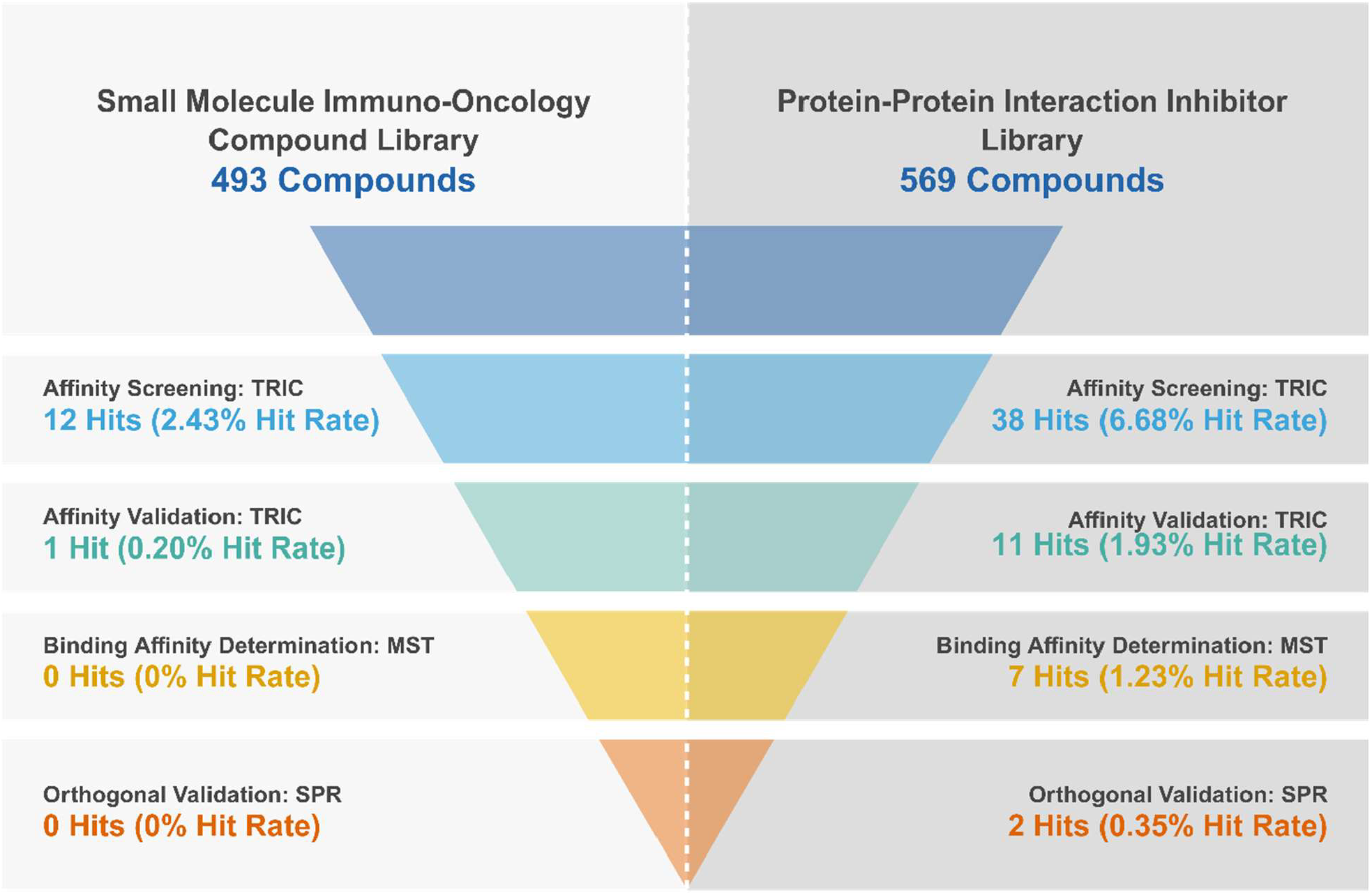
Funnel diagram summarizing the high-throughput screening and validation workflow for CD28-targeted small molecules. Two libraries: Small Molecule Immuno-Oncology Compound Library (MCE, 493 compounds) and Protein-Protein Interaction Inhibitor Library (MCE, 569 compounds); were used to perform the assay development. The multi-step method, comprising: Affinity Screening and Validation (TRIC), Binding Affinity Determination (MST), and Orthogonal Validation (SPR); shows the number of hits and the corresponding hit rates over the screening process. These results validate the primary screening assay’s capacity to identify and triage hits effectively, supporting its robustness and suitability for high-throughput applications targeting CD28 protein.

Sourced from the 10 mM stock plates, we prepared intermediate plates containing 2x concentrated small molecules (400 μM) in assay buffer supplemented with 4% DMSO. Two full columns were reserved as reference (negative control) wells, containing only assay buffer with the same DMSO concentration but no small molecules (**Figure 2**). The intermediate plates can be stored at -30°C for up to one week before use. Once CD28 protein is freshly labelled with the RED-tris-NTA dye, it is added to each sample within the 384-well plate and incubated for 15 minutes. After incubation, the plate was inserted into the Dianthus instrument and data acquisition was performed over a 35-minute reading time (**Figure 2**).

**Figure 2.**
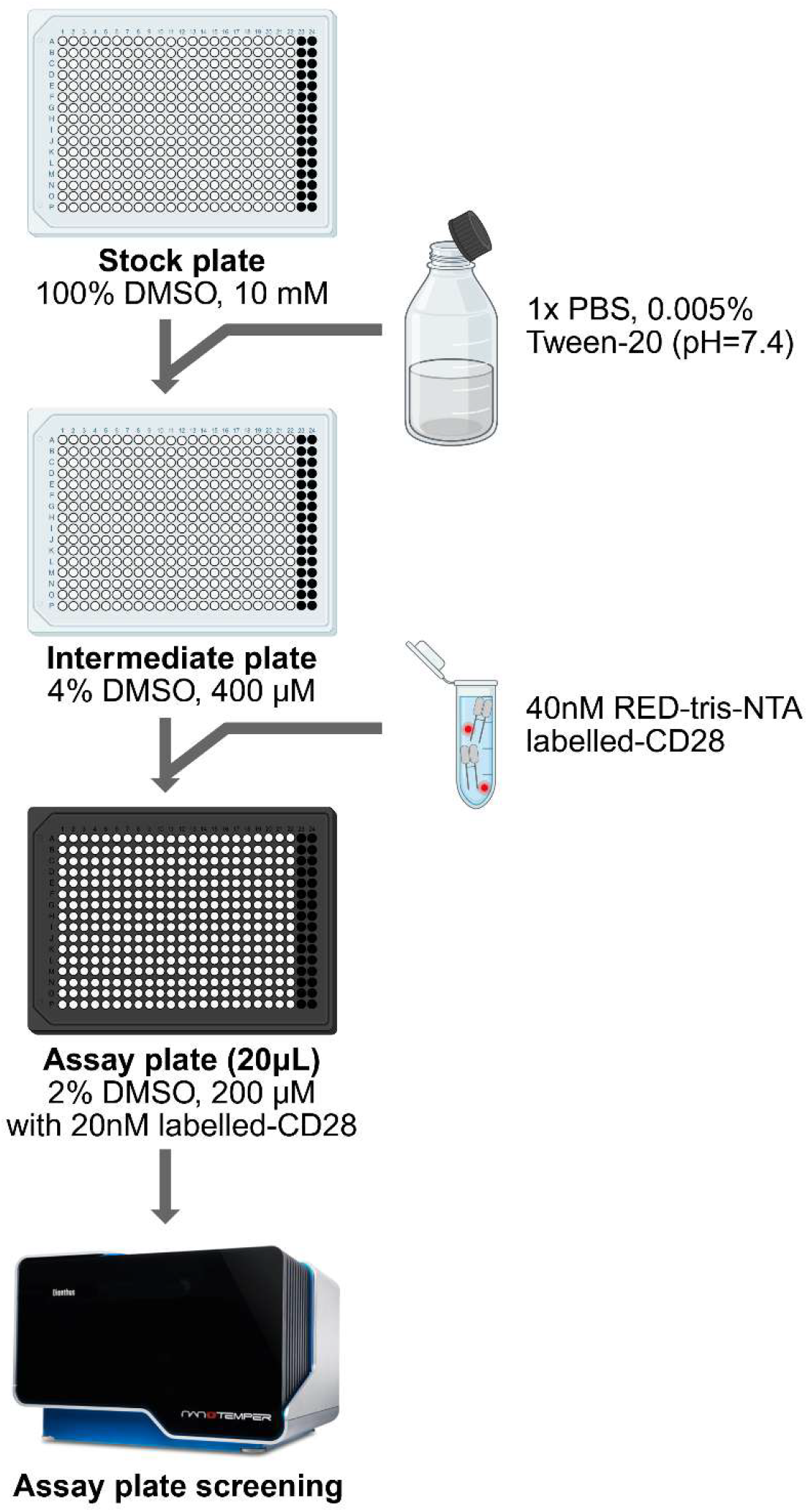
Workflow for the Preparation and Screening of Small Molecule Libraries Against RED-tris-NTA Labelled CD28. The process begins with a stock plate containing compounds in 100% DMSO at 10 mM, followed by dilution into an intermediate plate (4% DMSO, 400 μM). Final assay plates (Dianthus plates, 20 μL) are prepared with compounds at 200 μM and 20 nM of the labelled CD28 protein in 1x PBS with 0.005% Tween-20 supplemented with 2% DMSO (pH=7.4). Binding affinity screening of the small molecules to the target protein is performed using the Dianthus instrument (NanoTemper Technologies).

### 3.2. Data Analysis and Hit Identification from HTS Using TRIC Technology

Based on the TRIC technology, we obtained a single F_norm_ value for each screened sample, both small molecules and negative control wells, and plotted them (**Figures 3-4**). As previously described^8^, tested compounds that resulted in a response amplitude (ΔF_norm_) greater than three times the standard deviation of the negative control were initially considered CD28 binders. This threshold is indicated by black dashed lines in **Figures 3-4**. However, due to a higher-than-expected hit rate and the previously determined ΔF_norm_ value of the positive control CD80, we raised the threshold to a ΔF_norm_ of 10 units (approximately 50% of the positive control signal amplitude) for a compound to be considered a hit. This adjusted threshold is indicated by the grey shaded area in **Figures 3-4**. All compounds that showed any scan anomaly, autofluorescence, quenching effect towards the RED-tris-NTA dye, or aggregation at the tested concentration were excluded for subsequent experiments (**Figures 3-4**, red crosses). All remaining small molecules, represented as orange dots above or below the grey shaded areas in Figures 3 and 4, were cherry-picked from the stock plates and considered as potential hit compounds.

**Figure 3.**
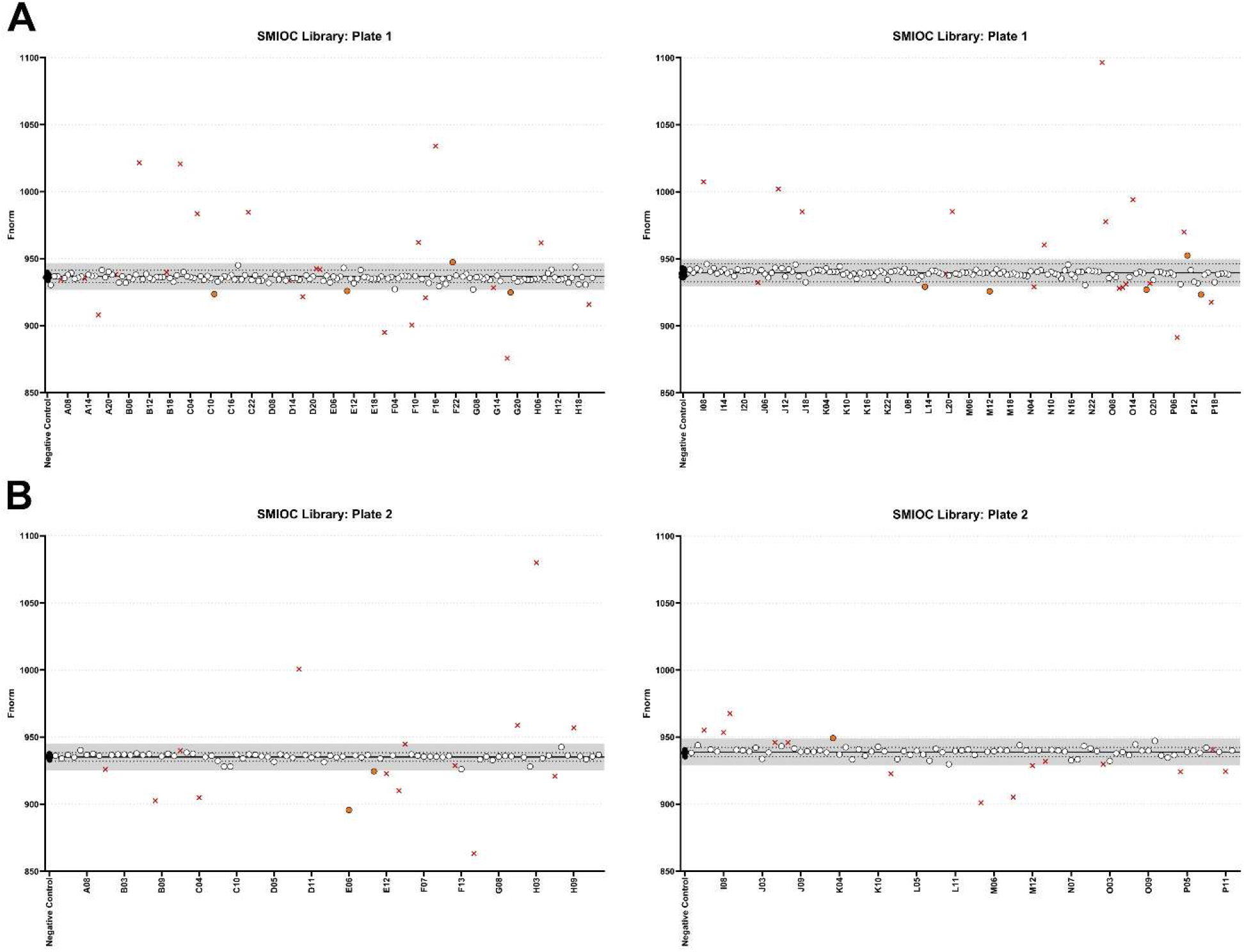
Identification of Hits in the Small Molecule Immuno-Oncology Compound (SMIOC) Library via HTS Using TRIC Technology. Each data point represents the normalized fluorescence value (F_norm_) obtained for an individual sample. Two sets of 16 negative control samples (black dots) were included on each half of the plate to reduce potential laser batch effects during data acquisition (as each half plate is read by a separate laser). The mean value of these controls (solid black line) and their standard deviation were used to set an initial threshold for identifying potential CD28 binders. The initial threshold was defined as a response amplitude (ΔF_norm_) exceeding three times the standard deviation of the negative control (black dashed lines), following the empirical rule. Due to a higher-than-expected hit rate and the known ΔF_norm_ value of the CD80 positive control, the threshold was subsequently raised to ΔF_norm_=10 units, corresponding to approximately 50% of the positive control signal amplitude. This adjusted threshold is indicated by the grey shaded area. Compounds exhibiting scan anomalies, autofluorescence, quenching effects toward the RED-tris-NTA dye, or aggregation at the tested concentration (200 μM) were excluded from further analysis and are marked with red crosses. The remaining 12 small molecules, shown as orange dots located above or below the grey shaded area, were identified as potential CD28 binders from the initial 493 compounds in the SMIOC library (MCE). The graphs show the results from a single HTS experiment using Dianthus instrument (NanoTemper Technologies).

**Figure 4.**
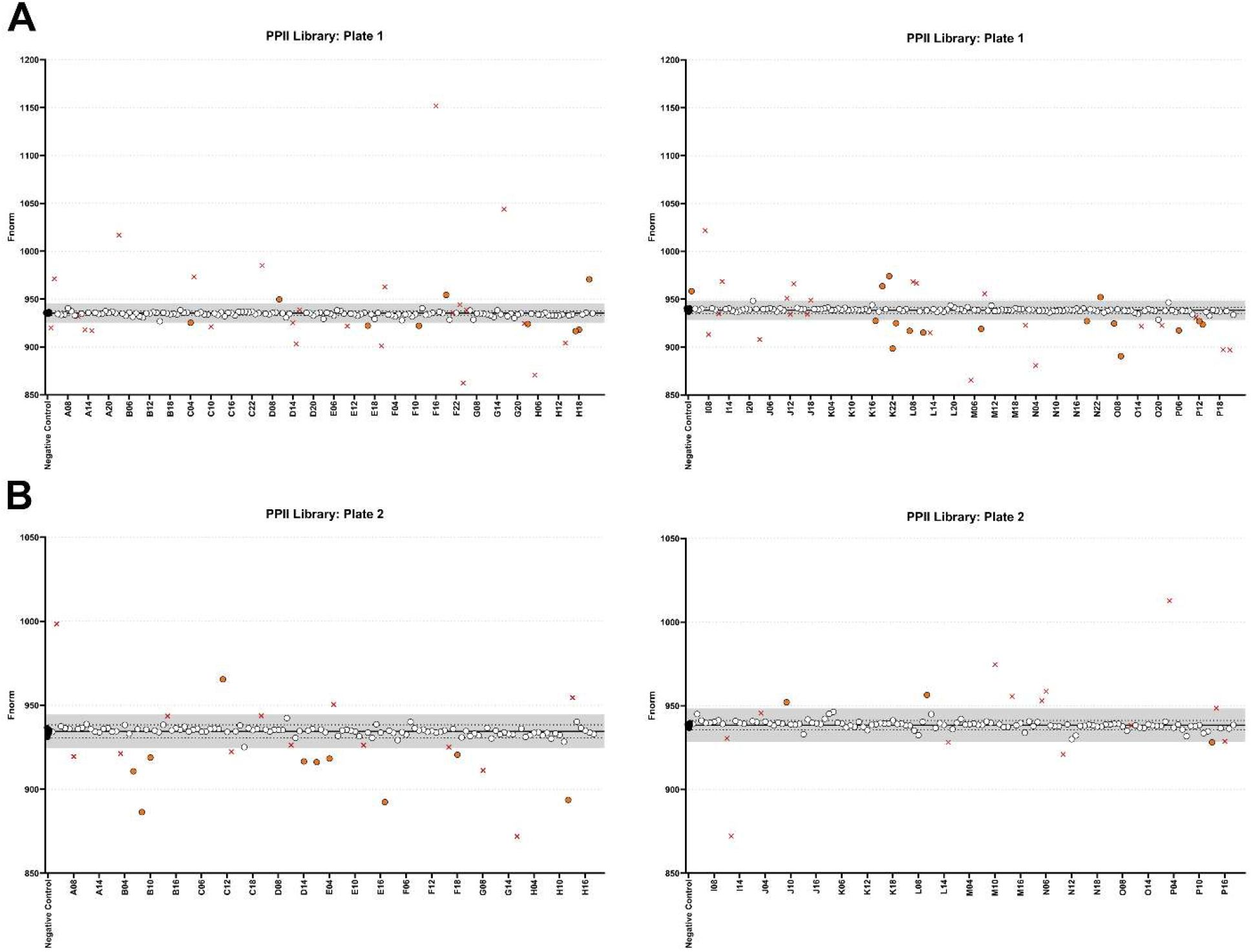
Identification of Hits in the Protein-Protein Interaction Inhibitor (PPII) Library via HTS Using TRIC Technology. This figure presents results analogous to those shown in Figure 3 but using a different compound library. Each data point represents the normalized fluorescence value (F_norm_) obtained for an individual sample. Two sets of 16 negative control samples (black dots) were included on each half of the plate to reduce potential laser batch effects during data acquisition (as each half plate is read by a separate laser). The mean value of these controls (solid black line) and their standard deviation were used to set an initial threshold for identifying potential CD28 binders. The initial threshold was defined as a response amplitude (ΔF_norm_) exceeding three times the standard deviation of the negative control (black dashed lines), following the empirical rule. Due to a higher-than-expected hit rate and the known ΔF_norm_ value of the CD80 positive control, the threshold was subsequently raised to ΔF_norm_=10 units, corresponding to approximately 50% of the positive control signal amplitude. This adjusted threshold is indicated by the grey shaded area. Compounds exhibiting scan anomalies, autofluorescence, quenching effects toward the RED-tris-NTA dye, or aggregation at the tested concentration (200 μM) were excluded from further analysis and are marked with red crosses. The remaining 38 small molecules, shown as orange dots located above or below the grey shaded area, were identified as potential CD28 binders from the initial 569 compounds in the PPII library (MCE). The graphs show the results from a single HTS experiment using the Dianthus instrument (NanoTemper Technologies).

Two additional experiments were then conducted under identical conditions to validate the ΔF_norm_ values (**Table 2**). Finally, an additional control experiment (*details in the Methods section*) was performed by incubating individual hits with the RED-tris-NTA dye alone. This allowed us to exclude some compounds due to interference with the dye itself, either showing an increase or decrease of the dye fluorescence intensity or a change in the TRIC trace in the absence of protein, when compared to the RED-tris-NTA dye alone in assay buffer supplemented with 2% DMSO (**Table 2**). Thus, after the affinity validation step, which included three independent binding check experiments as well as a control assay, one compound out of 493 initial compounds in the SMIOC library and 11 compounds out of 569 initial compounds in the PPII library were selected for binding affinity determination (**Table 3**).

**Table 2.** Screening Results of Small Molecules from the SMIOC and PPII Libraries Following Affinity Validation. Twelve compounds from the Small Molecule Immuno-Oncology Compound (SMIOC) library and 38 from the Protein-Protein Interaction Inhibitor (PPII) library were selected based on a single-dose (200 μM) high-throughput screening (HTS) experiment. These compounds were further evaluated at the same concentration across three independent replicates. The table presents the mean normalized fluorescence (F_norm_) values derived from these replicates. For each screening plate (divided into top and bottom halves, corresponding to each infrared laser in the instrument), a data range (ΔF_norm_) was defined for the negative control set, spanning from 10 F_norm_ units below to 10 units above the mean F_norm_ of the negative control replicates. Compounds with mean F_norm_ values falling outside this ΔF_norm_ range, and that did not trigger any experimental flags in a subsequent control experiment (*details in the Methods section*), were selected for further binding affinity studies.

| SMIOC Library |  |  |  |
| --- | --- | --- | --- |
| Sample | Fnorm | Flag? | Selected? |
| Negative Control | 927.3 947.3 |  |  |
| 1C11 | 955.1 |  | ✓ |
| 1E10 | 930.6 |  | ✗ |
| 1F21 | 945.3 | YES | ✗ |
| 1G18 | 926.3 | YES | ✗ |
| Negative Control | 928.6 948.6 |  |  |
| 1L13 | 923.1 | YES | ✗ |
| 1M12 | 935.7 |  | ✗ |
| 1O18 | 930.4 |  | ✗ |
| 1P10 | 961.3 | YES | ✗ |
| 1P14 | 918.3 | YES | ✗ |
| Negative Control | 926.5 946.5 |  |  |
| 2E06 | 917.9 | YES | ✗ |
| 2E10 | 926.8 |  | ✗ |
| Negative Control | 928.2 948.2 |  |  |
| 2K03 | 946 | YES | ✗ |

| PPII Library |  |  |  |
| --- | --- | --- | --- |
| Sample | Fnorm | Flag? | Selected? |
| Negative Control | 926.6 946.6 |  |  |
| 1C04 | 930.1 |  | ✗ |
| 1D10 | 959.5 |  | ✓ |
| 1E16 | 927.1 |  | ✗ |
| 1F11 | 916 | YES | ✗ |
| 1F19 | 958.5 |  | ✓ |
| 1H03 | 933.7 | YES | ✗ |
| 1H17 | 931.2 | YES | ✗ |
| 1H18 | 928 |  | ✗ |
| 1H21 | 969.2 |  | ✓ |
| Negative Control | 928 948 |  |  |
| 1I03 | 962 |  | ✓ |
| 1K17 | 929.4 | YES | ✗ |
| 1K19 | 957.8 |  | ✓ |
| 1K21 | 969.9 | YES | ✗ |
| 1K22 | 902.8 | YES | ✗ |
| 1L03 | 929.3 |  | ✗ |
| 1L07 | 913.7 | YES | ✗ |
| 1L11 | 930.6 | YES | ✗ |
| 1M08 | 919.1 | YES | ✗ |
| 1N19 | 938.9 | YES | ✗ |
| 1O03 | 951.8 |  | ✓ |
| 1O07 | 930.2 | YES | ✗ |
| 1O09 | 909 | YES | ✗ |
| 1P06 | 917.7 | YES | ✗ |
| 1P12 | 935.8 | YES | ✗ |
| 1P13 | 931.8 | YES | ✗ |
| Negative Control | 926.2 946.2 |  |  |
| 2B06 | 919 |  | ✓ |
| 2B08 | 889.4 |  | ✓ |
| 2B10 | 916 |  | ✓ |
| 2C11 | 959.9 |  | ✓ |
| 2D14 | 926.5 | YES | ✗ |
| 2D17 | 928.2 | YES | ✗ |
| 2E04 | 917.9 | YES | ✗ |
| 2E17 | 804.5 | YES | ✗ |
| 2F18 | 932.1 | YES | ✗ |
| 2H12 | 918 | YES | ✗ |
| Negative Control | 928 948 |  |  |
| 2J09 | 950 | YES | ✗ |
| 2L10 | 954.1 |  | ✓ |
| 2P13 | 928.9 |  | ✗ |

**Table 3.**
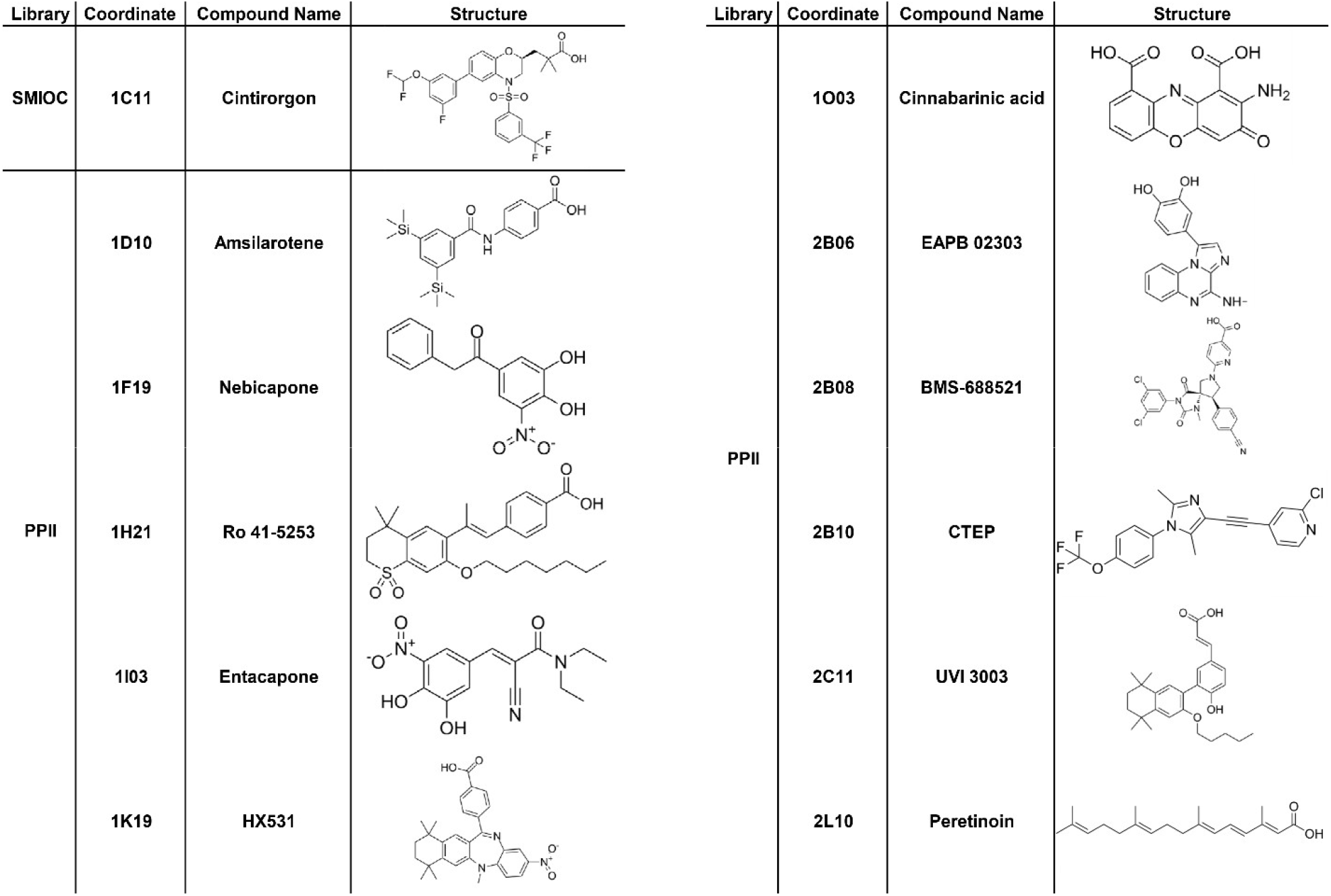
List of Selected Small Molecules from the SMIOC and PPII Libraries Following Affinity Validation. One compound from the Small Molecule Immuno-Oncology Compound (SMIOC) library and 11 from the Protein-Protein Interaction Inhibitor (PPII) library were selected based on the affinity validation step, which included three independent binding assays at 200 μM and a control experiment (*details provided in the Methods section*). For each selected compound, the table includes the library name, plate coordinate, compound name, and chemical structure.

Based on these results, we have successfully developed and optimized a primary screening assay that performs reliably at scale and effectively identifies hits. These findings demonstrate that the assay is both robust and suitable for HTS applications, providing proof of concept that HTS for the CD28 protein is feasible and effective using the current approach.

### 3.3. Binding Affinity Characterization of CD28 Hits Using MST

Our follow-up objective was to confirm whether a dose-response binding to the CD28 protein could be observed for each of the identified CD28 binders, and consequently, whether a K_d_ could be determined for each compound. To achieve this, we employed the same technology (TRIC) but used a different instrument, the Monolith (NanoTemper Technologies), to determine the K_d_ values. We successfully determined K_d_ values in the micromolar range for seven compounds, all of which originated from the PPII library (**Figure 5**). In contrast, the remaining five compounds: Cintirorgon, Amsilarotene, Ro 41-5253, BMS-688521, and UVI 3003; did not exhibit a dose-dependent response or yielded inconsistent K_d_ values across replicates.

**Figure 5.**
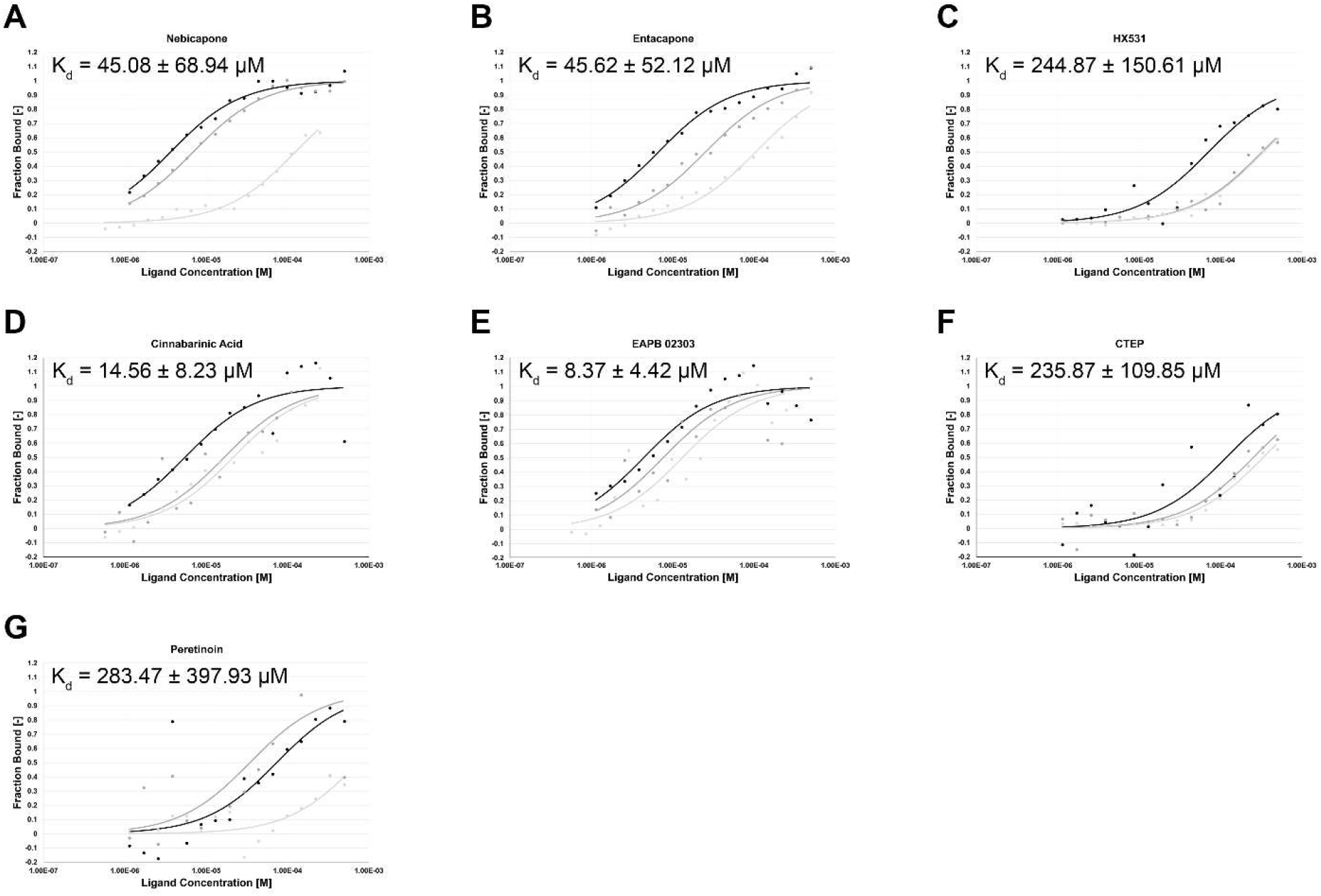
Dose-Response Binding Curves for Seven CD28-Binding Compounds from the PPII Library. Binding affinities were measured using TRIC technology on the Monolith instrument (NanoTemper Technologies). Each graph displays the fraction bound as a function of ligand concentration (logarithmic scale) for one compound. Sixteen-point serial dilutions were prepared for each candidate small molecule in assay buffer supplemented with 2% DMSO (ranging from 500 μM to 571 nM). K_d_ values were derived from three independent experiments using the K_d_ fit model, which describes a 1:1 binding interaction according to the law of mass action. All compounds exhibited binding in the micromolar range. The MST-on time for all compounds (Nebicapone, Entacapone, HX531, Cinnabarinic Acid, EAPB 02303, CTEP, and Peretinoin) was 5 seconds.

### 3.4. Orthogonal Validation of Small-Molecule CD28 Binders Using SPR

SPR is a label-free technique that allows direct measurement of small molecule binding to target proteins. Its high sensitivity and real-time detection capabilities make it a strong complementary approach to TRIC-based assays. An orthogonal validation using SPR confirmed that both EABP 02303 (518.67 ± 286.23 μM) and CTEP (181.50 ± 53.03 μM) small molecules bind to human CD28 protein and that their binding affinities are in the micromolar range (**Figure 6**).

**Figure 6.**
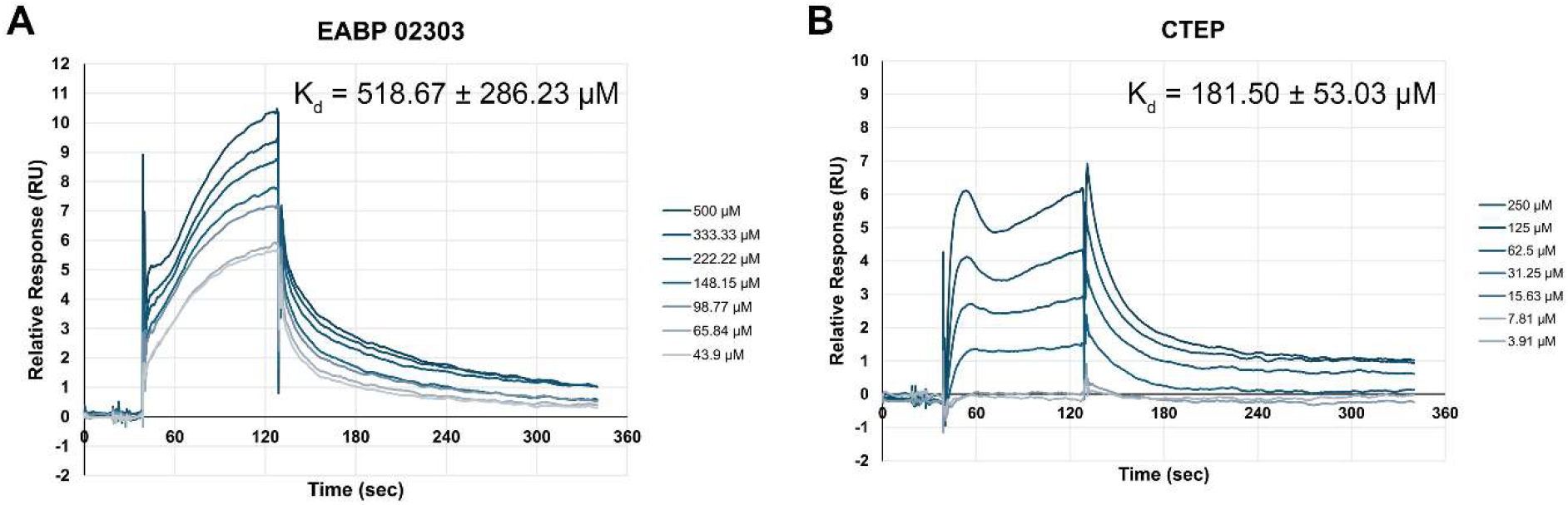
Validation of Two Small-Molecule CD28 Binders via SPR Analysis. **A**. Serial dilutions of the EABP 02303 compound (ranging from 500 to 43.9 μM, including a 0 μM reference; 1.5-fold dilution series) were injected onto a Sensor Chip CAP with immobilized human CD28 protein using a multi-cycle kinetics method. **B**. Serial dilutions of the CTEP compound (ranging from 250 to 3.91 μM, including a 0 μM reference; 2-fold dilution series) were similarly injected onto a Sensor Chip CAP with immobilized human CD28 protein using a multi-cycle kinetics method. The data represent the relative response units (RU) over time during the 90-second association phase and the 240-second dissociation phase of a representative experiment. Binding curves were analyzed using non-linear curve fitting based on steady-state affinity analysis.

In summary, we have successfully developed and validated a robust TRIC-based HTS assay for identifying small molecule binders of human CD28. This scalable, immobilization-free platform represents a significant advancement in the field of immunomodulatory drug discovery, enabling rapid assessment of chemical libraries against a challenging target. As a proof of concept, our screening campaign yielded multiple hits, several of which demonstrated micromolar binding affinity and were further substantiated through orthogonal biophysical techniques including MST and SPR.

Importantly, this is the first report to demonstrate the application of TRIC technology for CD28 target engagement, establishing its feasibility and reliability in early-stage screening. By integrating orthogonal validation and rigorous artifact exclusion, we have increased the confidence in our primary hits and built a solid foundation for future structure–activity relationship (SAR) studies and hit-to-lead optimization.

Looking forward, our TRIC platform provides a valuable tool for expanding small-molecule approaches to CD28 modulation and opens new avenues for dual-target therapeutic strategies involving CD28 and ICOS in inflammatory and autoimmune diseases. These findings underscore the potential of TRIC-enabled screening as a generalizable approach for discovering modulators of other immune checkpoint receptors.

## CRediT Authorship Contribution Statement

Laura Calvo-Barreiro: Conceptualization, Methodology, Validation, Formal Analysis, Investigation, Data Curation, Writing – Original Draft, Visualization. Moustafa T. Gabr: Writing - Review & Editing, Funding Acquisition.

## Declaration of Competing Interests

LC-B and MTG declare no competing financial interests.

## Acknowledgments

This work was supported by the National Institute of Diabetes and Digestive and Kidney Diseases (NIDDK) under grant number R01DK137299. We would like to thank the Fisher Drug Discovery Resource Center of Rockefeller University (RRID:SCR_020985) for providing access to the Nanotemper Dianthus NT.23 Pico, Nanotemper Monolith NT.115, and Cytiva Biacore 8K instruments.

## Data Availability Statement

The data that support the findings of this study are available in the Supporting Information.

## Notes

### Competing Interest Statement

The authors have declared no competing interest.

