## Supporting Information for "TRIC-Based High-Throughput Screening Enables the Discovery of Small Molecule CD28 Binders"

### 1. Chromatographic and Mass Spectrum Data for Selected Small Molecule from the Small Molecule Immuno-Oncology Compound (SMIOC) Library

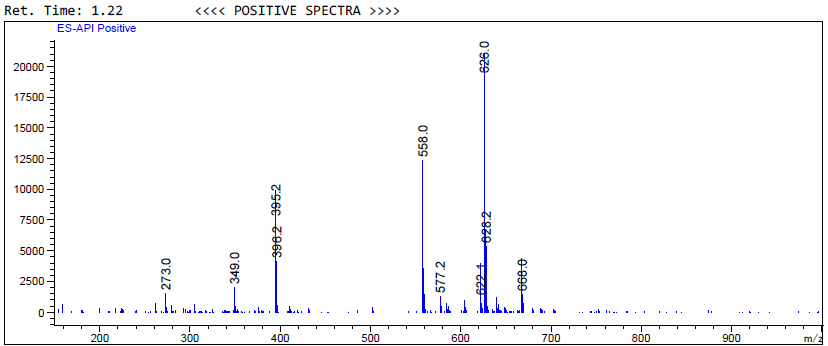

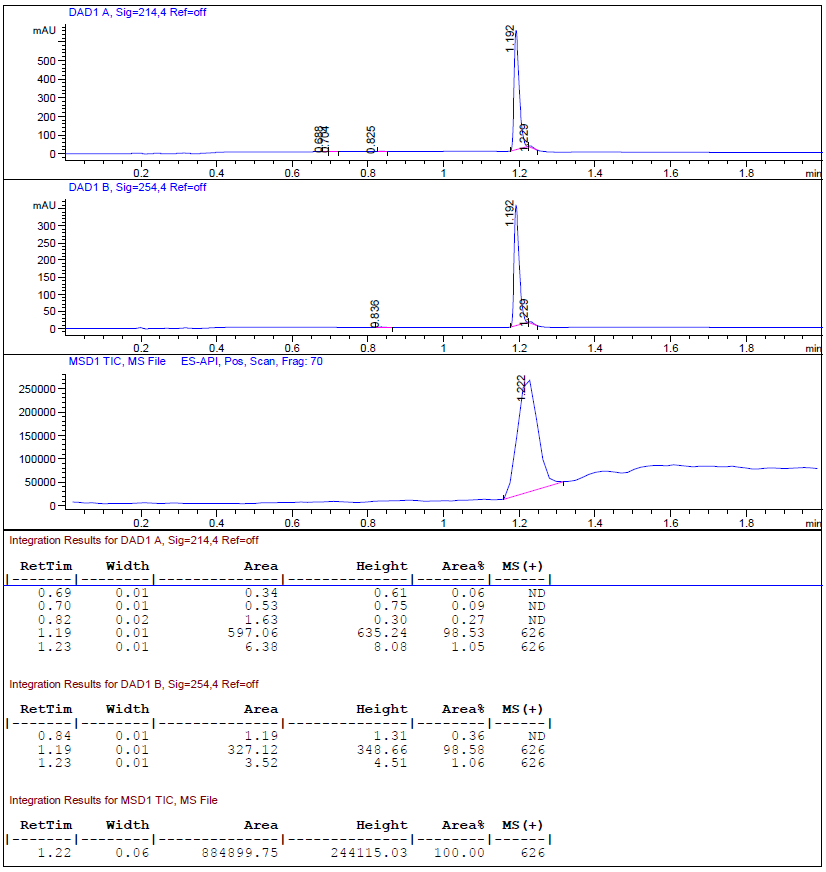

**Figure S1.** Chromatographic data and mass spectrum from UPLC-MS analysis of Cintirorgon.

### 2.  Chromatographic and Mass Spectrum Data for Selected Small Molecules from the Protein-Protein Interaction Inhibitor (PPII) Library

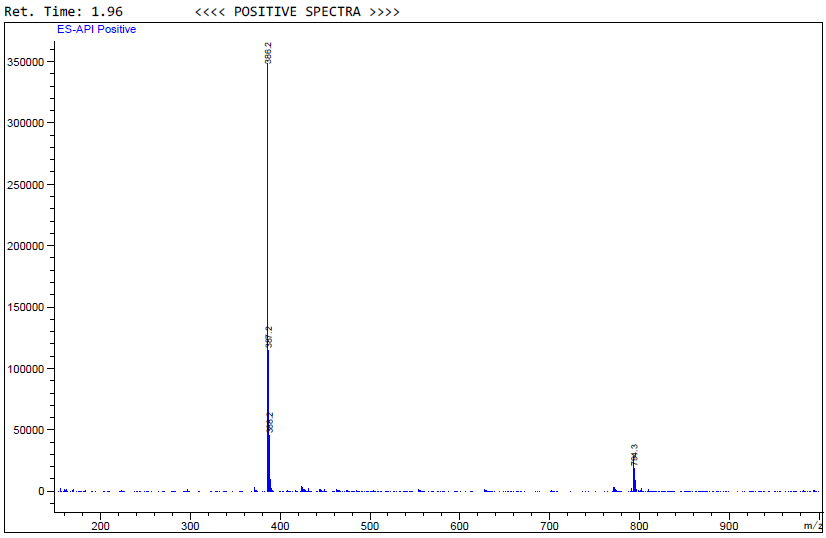

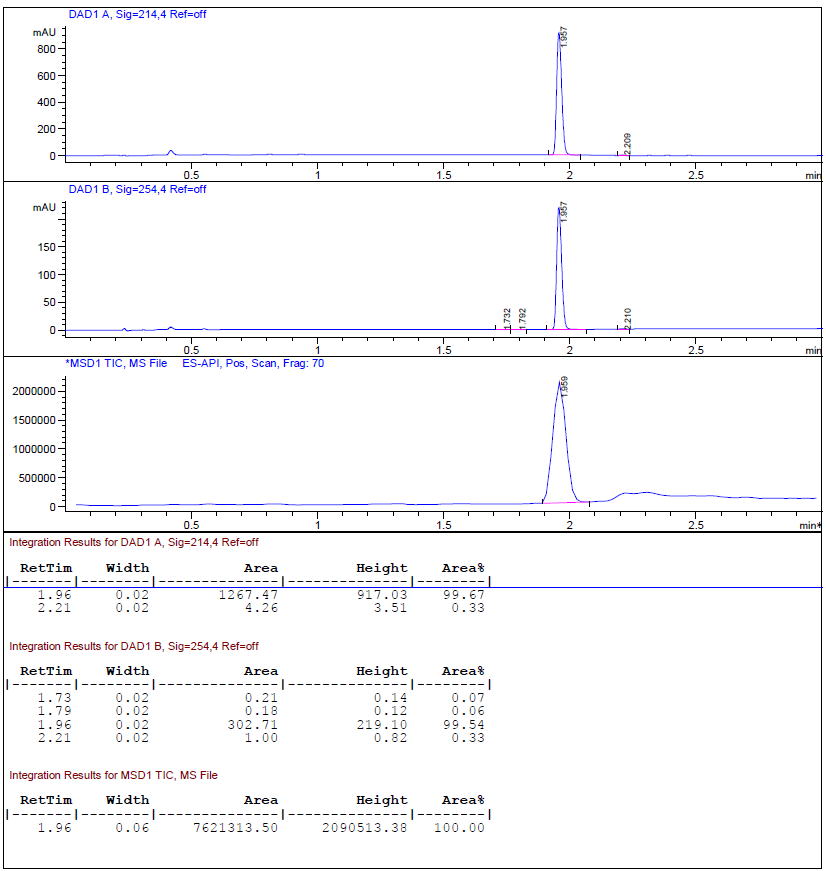

**Figure S2.** Chromatographic data and mass spectrum from UPLC-MS analysis of Amsilarotene.

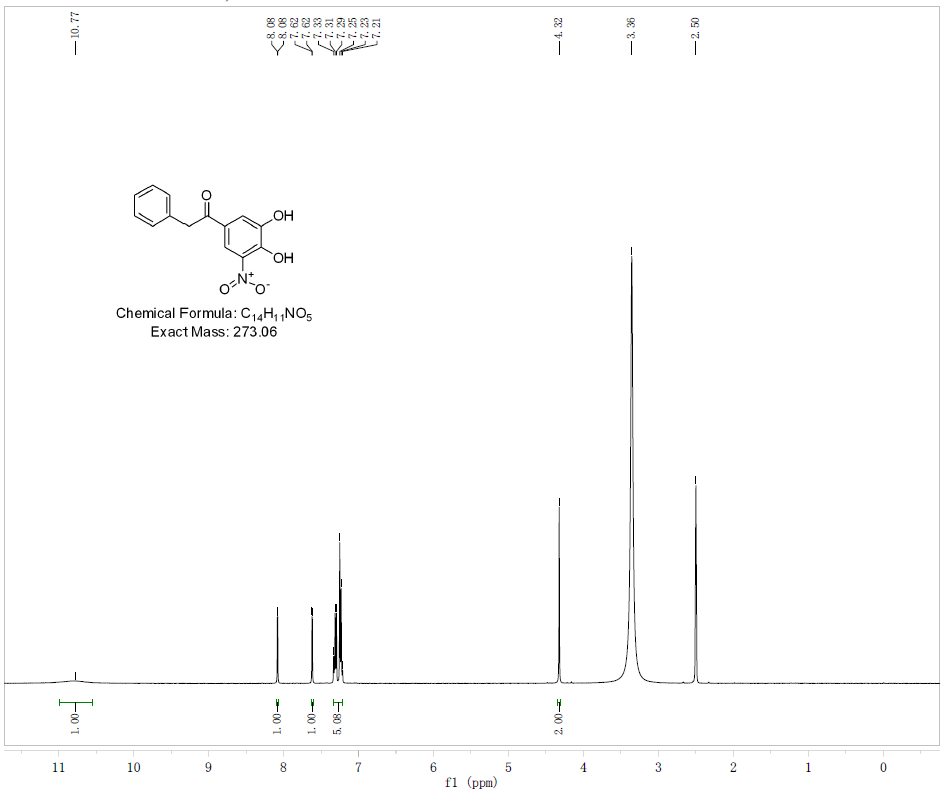

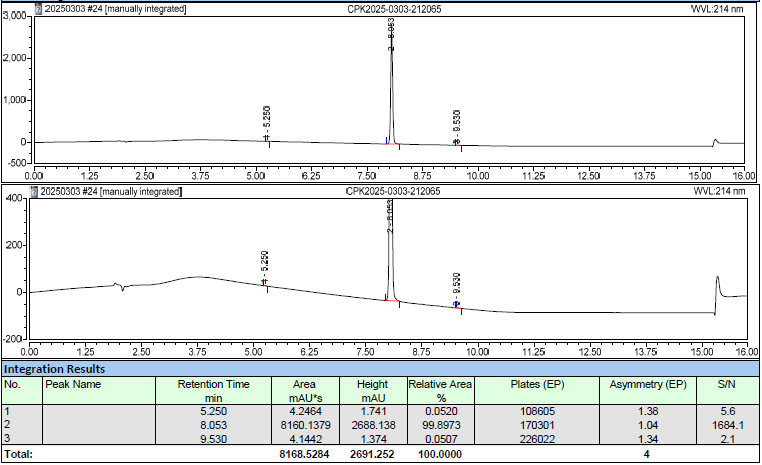

**Figure S3.** Chromatographic data and mass spectrum from UPLC-MS analysis of Nebicapone.

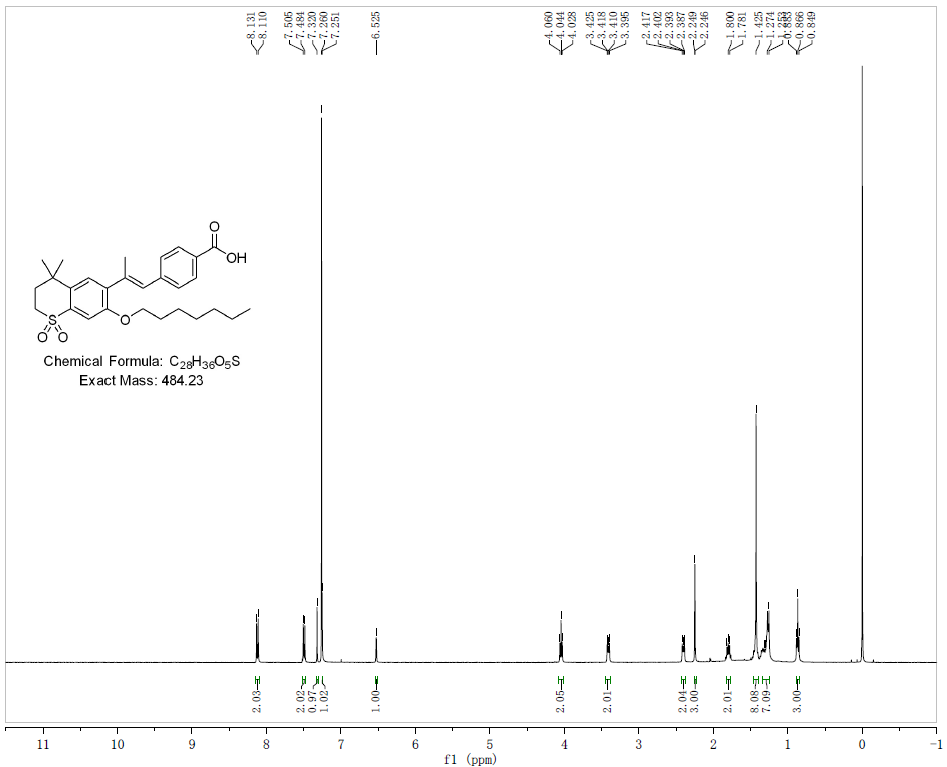

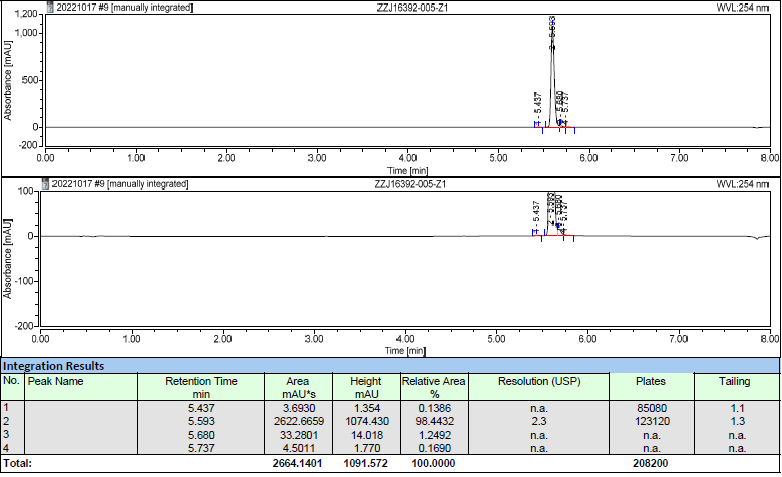

**Figure S4.** Chromatographic data and mass spectrum from UPLC-MS analysis of Ro 41-5253.

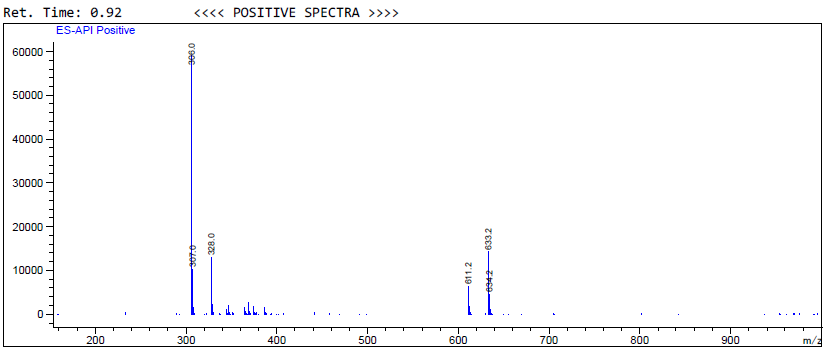

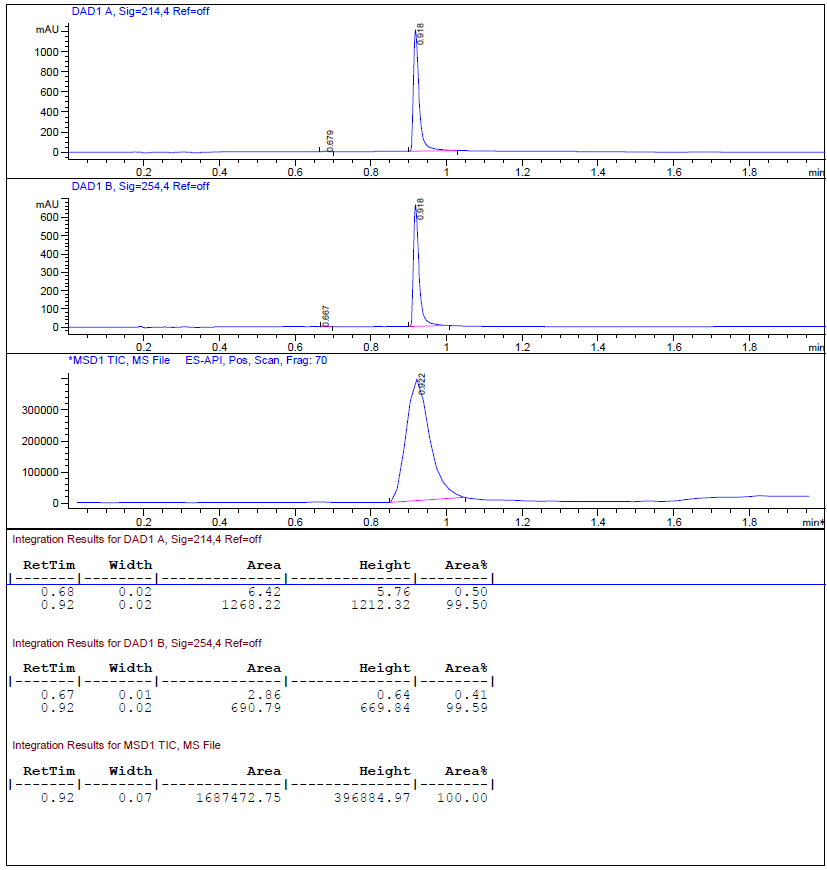

**Figure S5.** Chromatographic data and mass spectrum from UPLC-MS analysis of Entacapone.

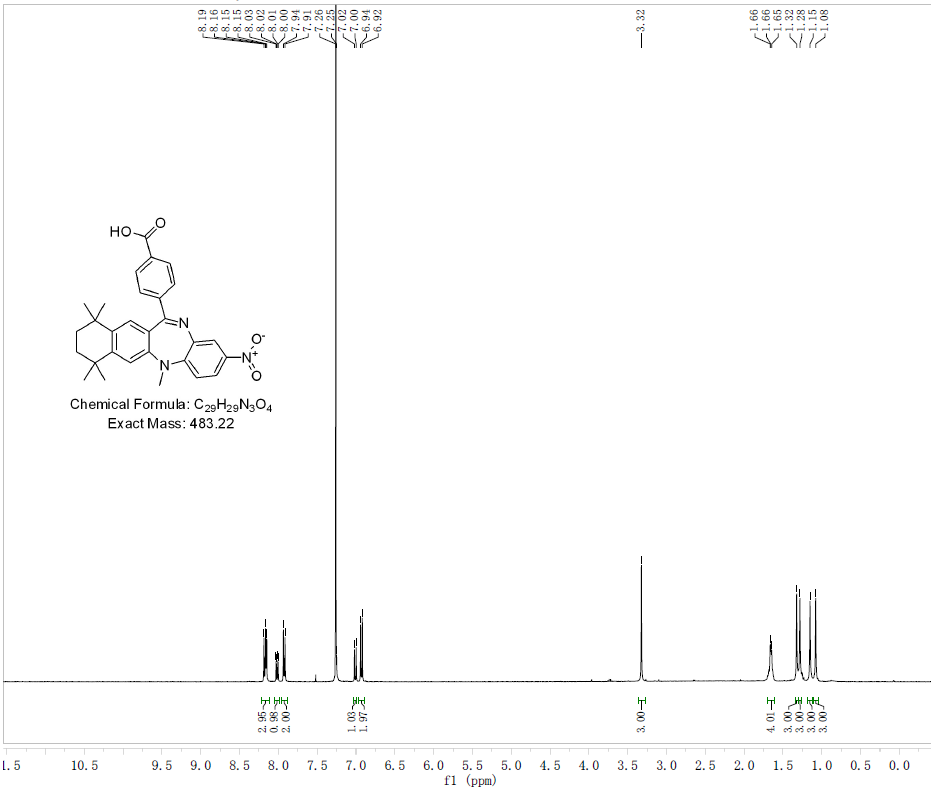

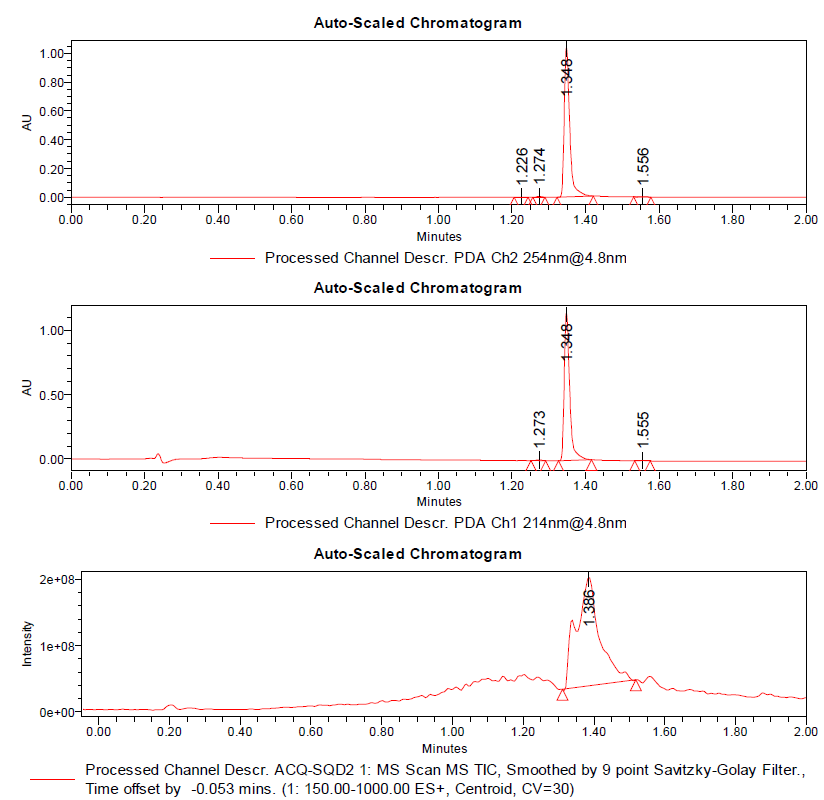

**Figure S6.** Chromatographic data and mass spectrum from UPLC-MS analysis of HX531.

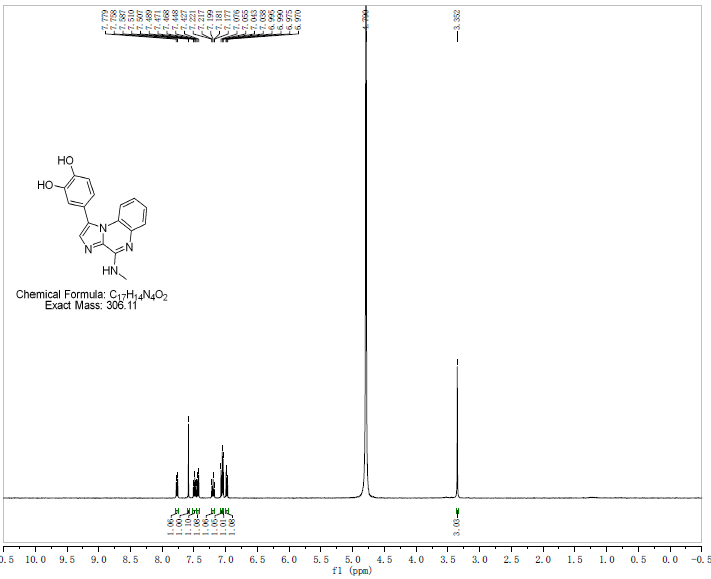

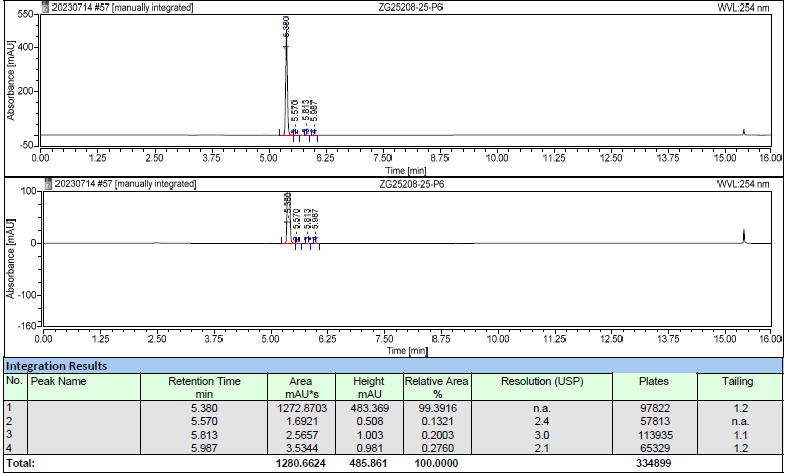

**Figure S7.** Chromatographic data and mass spectrum from UPLC-MS analysis of EAPB 02303.

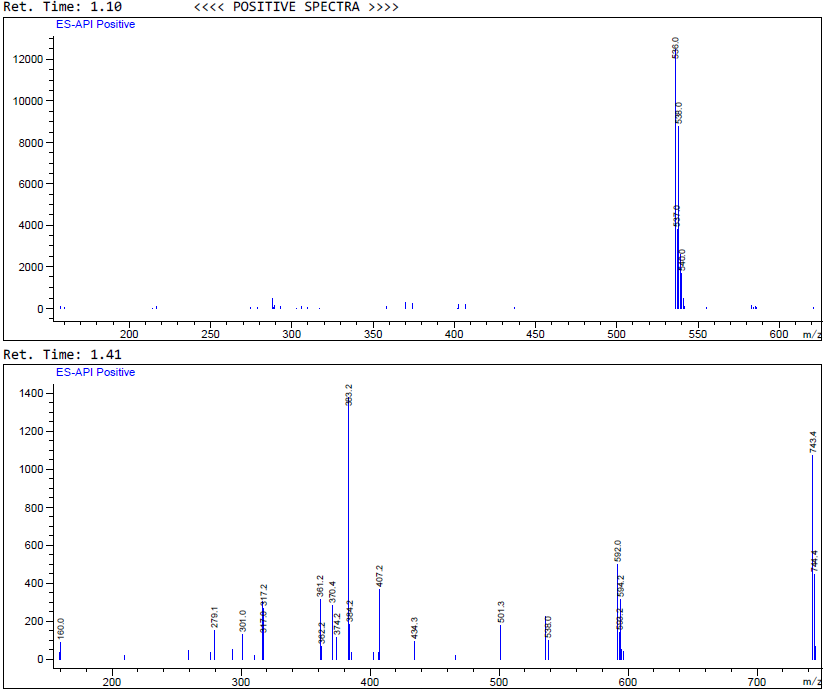

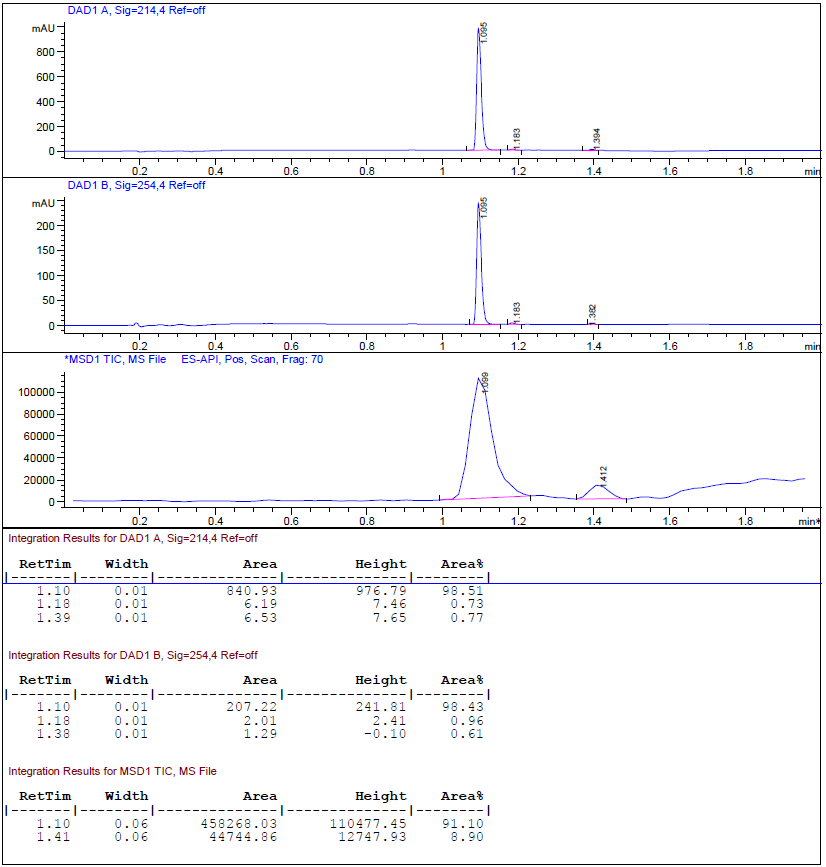

**Figure S8.** Chromatographic data and mass spectrum from UPLC-MS analysis of BMS-688521.

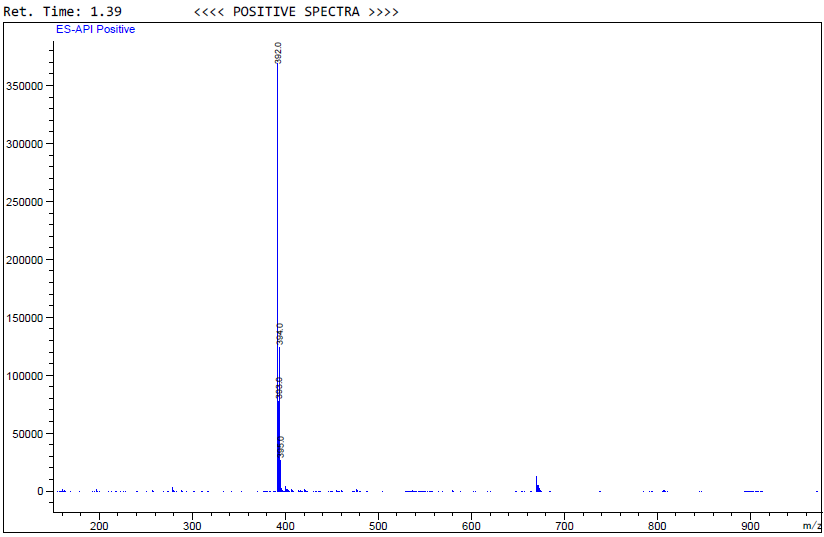

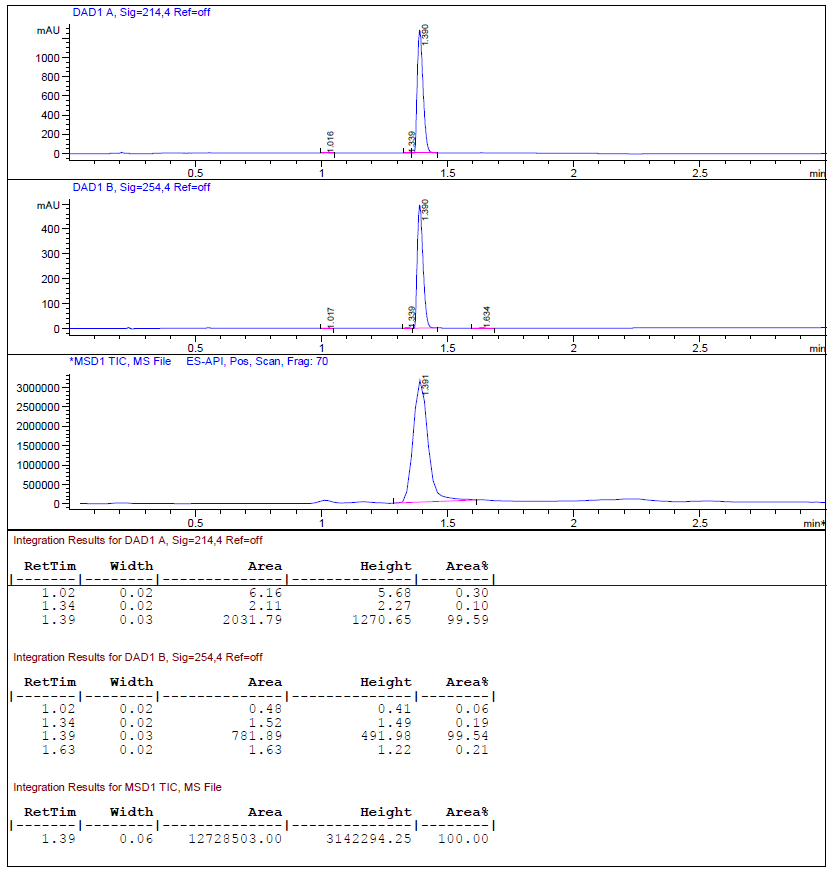

**Figure S9.** Chromatographic data and mass spectrum from UPLC-MS analysis of CTEP.

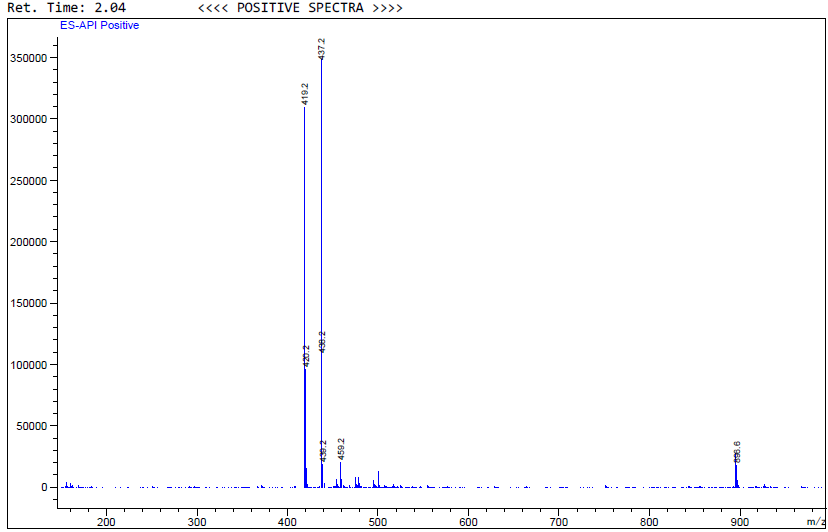

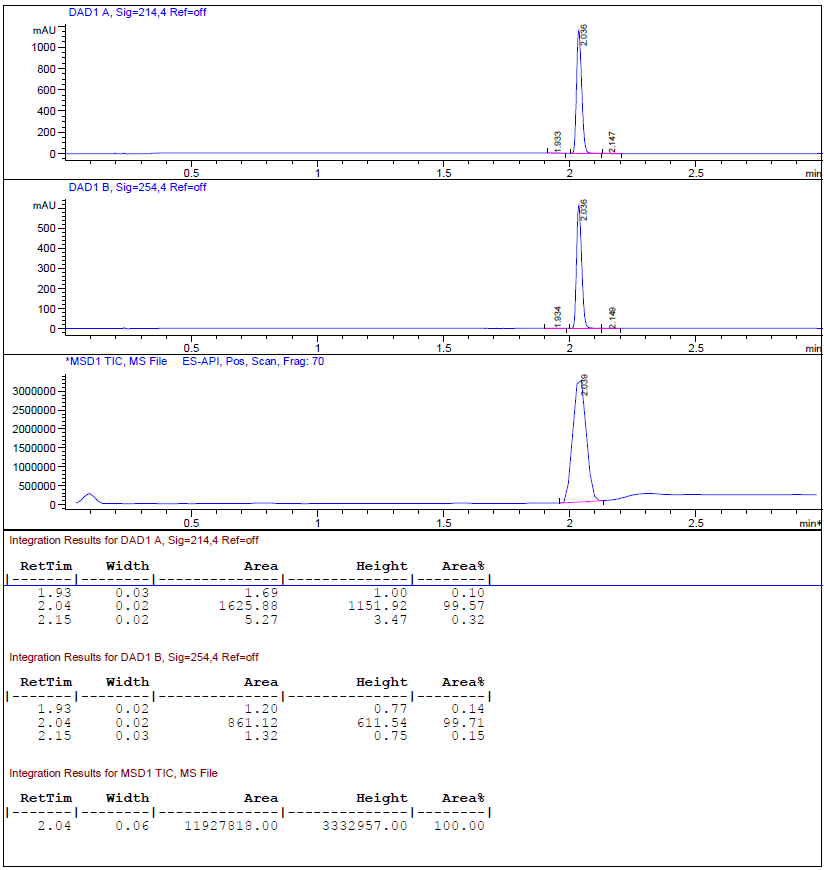

**Figure S10.** Chromatographic data and mass spectrum from UPLC-MS analysis of UVI 3003.

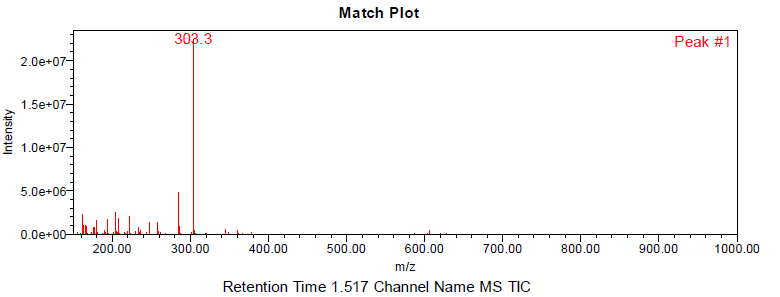

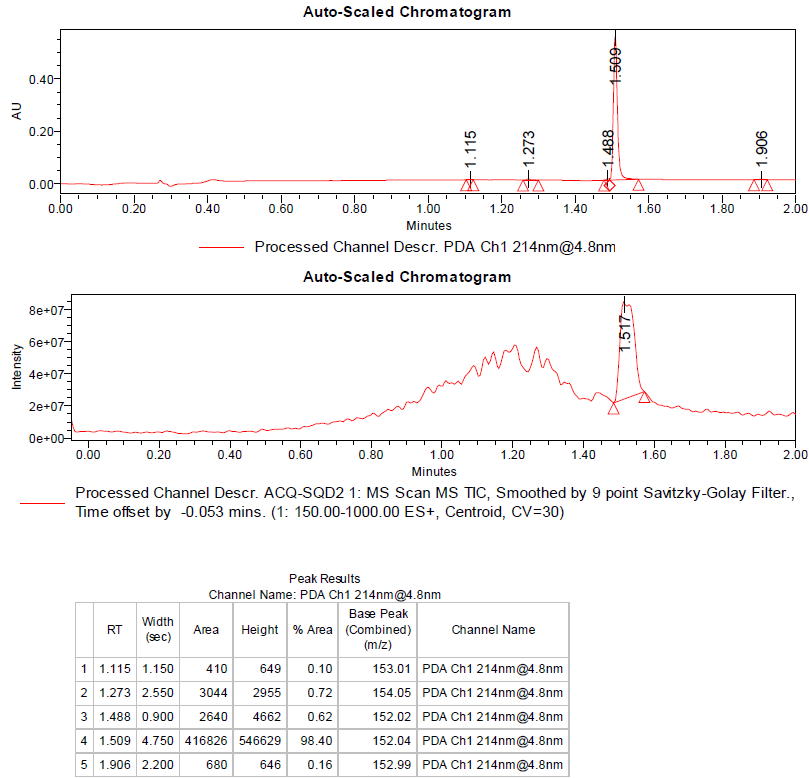

**Figure S11.** Chromatographic data and mass spectrum from UPLC-MS analysis of Peretinoin.
